# The novel reversible LSD1 inhibitor SP-2577 promotes anti-tumor immunity in SWItch/Sucrose-NonFermentable (SWI/SNF) complex mutated ovarian cancer

**DOI:** 10.1101/2020.01.10.902528

**Authors:** Raffaella Soldi, Tithi Ghosh Halder, Alexis Weston, Trason Thode, Kevin Drenner, Rhonda Lewis, Mohan R. Kaadige, Shreyesi Srivastava, Sherin Daniel Ampanattu, Ryan Rodriguez del Villar, Jessica Lang, Hariprasad Vankayalapati, Bernard Weissman, Jeffrey M. Trent, William P.D. Hendricks, Sunil Sharma

## Abstract

Chromatin remodeling SWItch/Sucrose-NonFermentable (SWI/SNF) complexes, initially identified in yeast 20 years ago, are evolutionarily conserved multi-subunit protein complexes that use the energy from hydrolysis of adenosine triphosphate (ATP) to remodel nucleosome structure and modulate transcription. Mutations in proteins of SWI/SNF complexes occur in 20% of human cancers including ovarian cancer (OC). Approximately 50% of ovarian clear cell carcinoma (OCCC) carries mutations in the SWI/SNF subunit ARID1A while small cell carcinoma of the ovary hypercalcemic type (SCCOHT) is driven primarily by genetic inactivation of the SWI/SNF ATPase SMARCA4 (BRG1) alongside epigenetic silencing of the homolog ATPase SMARCA2 (BRM). Dual loss of these ATPases disrupts SWI/SNF chromatin remodeling activity and may also interfere with the function of other histone-modifying enzymes that associate with or are dependent on SWI/SNF activity. One such enzyme is lysine-specific histone demethylase 1 (LSD1/KDM1A) which regulates the chromatin landscape and gene expression by demethylating proteins, including histone H3. LSD1 associates with epigenetic complexes such as the nucleosome remodeling deacetylase complex (NuRD) and SWI/SNF to inhibit the transcription of genes involved in tumor suppression and cell differentiation. TGCA analysis of human cancers shows that LSD1 is highly expressed in SWI/SNF-mutated tumors. Further, SCCOHT and OCCC cell lines show low nM IC_50_s for the reversible LSD1 inhibitor SP-2577 (Seclidemstat, currently in clinical phase I trials), supporting that these SWI/SNF-deficient ovarian cancers are dependent on LSD1 activity. Recently, it has been also shown that inhibition of LSD1 stimulates interferon (IFN)-dependent anti-tumor immunity through induction of Endogenous Retroviruses Elements (ERVs) and may thereby overcome resistance to checkpoint blockade. Additionally, SCCOHTs have been shown to exhibit an immune-active tumor microenvironment with PD-L1 expression in both tumor and stromal cells that strongly correlated with T cell infiltration. Thus, in this study we investigated the ability of SP-2577 to promote anti-tumor immunity and T cell infiltration in SWI/SNF-mutant SCCOHT and OCCC models. Our data shows that the reversible LSD1 inhibitor SP-2577 stimulates IFN-dependent anti-tumor immunity in SCCOHT cells *in vitro* in a 3D immune-organoid platform. Additionally, SP-2577 promoted the expression of PD-L1 in both SCCOHT and OCCC models. Together our findings suggest that SP-2577 and checkpoint inhibitors as a therapeutic combination may induce or augment immunogenic responses in these tumors.

## Introduction

An increasing number of cancers are recognized to be driven partly by inactivation of subunits of the SWItch/Sucrose-NonFermentable (SWI/SNF) complex, a multi-protein ATP-dependent chromatin-remodeling complex with central roles in cell differentiation programs (1, 2). Pathogenic SWI/SNF mutations occur across diverse adult cancers, typically in a genomic background of numerous other driver mutations and/or genomic instability (3, 4). However, SWI/SNF driver mutations also occur in a unique subset of more uniform cancers, such as small cell carcinoma of the ovary hypercalcemic type (SCCOHT) (5), rhabdoid tumors (RT) (6, 7), thoracic sarcomas (8, 9), and renal medullary cancers (10). These cancers share genetic and phenotypic features even though they arise from different anatomic sites (1). Shared features include poorly differentiated morphology, occurrence in young populations, and clinically aggressive behavior (11, 12). Their genetic makeup is relatively simple, with an overall low tumor mutation burden, few structural defects, and, in most cases, universal inactivation of a single subunit in the SWI/SNF complex. Particularly, in ovarian cancers (OCs), the most lethal gynecologic malignancies in the developed world and the fifth leading cause of cancer-associated mortality among women in the United States (13), SWI/SNF alterations vary in different histologic subtypes. The ARID1A (BAF250a) subunit is mutated in approximately 50% of ovarian clear cell carcinomas (OCCC) and 30% of ovarian endometrioid carcinomas (OEC) (14). SCCOHT (15), a rare and very aggressive OC, is a single-gene disease with inactivating mutations in the subunit SMARCA4 (BRG1) (16–18) and epigenetic silencing of SMARCA2 (BRM) expression (17). SCCOHT is the most common undifferentiated ovarian malignant tumor in women under 40 years. In contrast, OCCC targets women aged 55 years or older and is characterized by mutations in phosphatidylinositol-4, 5-bisphosphate 3-kinase catalytic subunit α (PIK3CA) (19, 20) and phosphatase and tensin homolog (PTEN), in addition to the ARID1A mutations. Both SCCOHT and OCCC respond poorly to conventional chemotherapy, and to date there is no consensus on the optimal therapeutic strategy (5, 20–23).

ATP-dependent chromatin remodeling plays a critical role in cell differentiation through control of transcriptional programs. When disrupted, these programs result in abnormal gene expression that creates therapeutically targetable oncogenic dependencies (24). For example, in BRG1-deficient non-small cell lung cancers, BRM has been identified as a candidate synthetic lethal target (25, 26). Similarly, in BRG1-deficient small cell lung cancer, MYC-associated factor X (MAX) was identified as a synthetic lethal target (27). In ARID1A-mutated OC, inhibition of DNA repair proteins PARP and ATR, and the epigenetic factors EZH2, HDAC2, HDAC6 and BRD2 have all shown therapeutic promise (28). In SCCOHT, therapeutic vulnerabilities to receptor tyrosine kinase inhibitors (29), EZH2 inhibitors (30–32), HDAC inhibitors (33), bromodomain inhibitors (34), and CDK4/6 inhibitors (35, 36) have also been identified. Importantly, correlations between SWI/SNF mutations and responses to immune checkpoint inhibitors have also been observed (37). In renal cell carcinoma, patients carrying mutations in bromodomain-containing genes (PBRM1 and BRD8) showed exceptional response to the anti-CTLA-4 antibody Ipilimumab (38). A CRISPR screen to identify genes involved in anti-PD-1 resistance identified three SWI/SNF complex members as important determinants in melanoma (39). A moderate response to anti-PD-1 treatment was also reported in a cohort of four SCCOHT patients expressing PD-L1 (40), suggesting that the low tumor mutation burden is not a limitation for checkpoint immunotherapy. OCCC models have also recently been described to be responsive to checkpoint inhibition in combination with HDAC6 inhibitors, particularly in the ARID1A-deficient setting (41). In a Phase II clinical trial testing Nivolumab in platinum-refractory ovarian cancers, one of the two OCCC patients demonstrated a complete response (42). These data suggest that novel treatment approaches and combinations should be adopted to develop targeted therapies against SWI/SNF-mutant ovarian cancers.

LSD1 is an epigenetic enzyme that can either repress target gene expression by demethylating mono- or di-methylated histone H3 lysine 4 (H3K4me1/2) or activate targets by removing repressive H3K9me1/2. LSD1 is implicated in tumorigenesis and progression of many cancers and high LSD1 levels frequently correlate with aggressive cancer features (43–45). LSD1 also promotes tumor progression through demethylation of non-histone substrates such as p53, E2F1, DNMT1, and MYPT1, a regulator of RB1 phosphorylation (46–50). Further, recent studies indicate that LSD1 ablation can trigger anti-tumor immunity through stimulation of expression of endogenous retroviral elements (ERVs) and downregulation of expression of the RNA-induced silencing complex (RISC). The accumulation of double-stranded RNA (dsRNA) results in the stimulation of interferon (IFN) **β**-dependent immunogenic responses (51). These studies also show that LSD1 inhibition overcomes resistance to checkpoint blockade therapy *in vivo* by increasing tumor immunogenicity and T cell infiltration (51–53). Several studies have shown interaction between LSD1 and SWI/SNF complexes. In glioma, LSD1 is part of a co-repressor complex containing TLX, RCOR2 and the SWI/SNF core complex. Together, this co-repressor complex regulates stem-like properties of glioma initiating cells (GICs) (54). When associated with SWI/SNF, CoREST, HDAC1/2 and DMNTs, LSD1 regulates gene expression in the neural network underlying neurodegenerative diseases and brain tumors (55). In breast cancer, LSD1 associates with the SWI/SNF subunit SMARCA4 to form a hormone-dependent transcriptional repressor complex (56). A similar association is required for endogenous Notch-target gene expression in T-ALL cells (57). These findings suggest LSD1 as an important therapeutic target in cancers driven by SWI/SNF mutations.

In this study, we explored the therapeutic potential of SP-2577 (Seclidemstat, Salarius Pharmaceuticals, Houston TX), a potent reversible LSD1 inhibitor currently in Phase I clinical trials for Ewing Sarcoma (NCT03600649) and for advanced solid tumors (NCT03895684), to promote anti-tumor immunity and T cell infiltration in SWI/SNF-mutant OC. Our findings show that SP-2577 promotes ERV expression, activates the dsRNA-induced IFN pathway, and enhances T cell infiltration in SCCOHT. Treatment with SP-2577 also promotes PD-L1 expression, and thus can potentially overcome resistance to anti-PD-1 therapy in SCCOHT. Finally, the efficacy of SP-2577 to promote T cell infiltration is not exclusive to SCCOHT. We observed similar effects in other SWI/SNF mutation-dependent OCs, such as ARID1A-mutant OCCC. Our data strongly suggest LSD1 as a potential therapeutic target in SCCOHT and provides preclinical evidence supporting the combinatorial use of SP-2577 with anti-PD-L1 for SWI/SNF-mutation-dependent OCs.

## Materials and Methods

### Cell Lines and 2D Culture Maintenance

SCCOHT cell lines BIN67 and SCCOHT-1 (Generously donated by Dr. William Hendrick, TGen) were cultured in RPMI Medium 1640 with L-Glutamine (Gibco) and supplemented with 10% FBS (Gibco) and 1% penicillin/streptomycin (Gibco). SCCOHT cell line COV434 (Generously donated by Dr. William Hendricks, TGen), OCCC cell line TOV21G (ATCC), and the doxycycline-inducible COV434 pIND20 BRG1-2.7 and TOV21G pIND20 ARID1A (generously donated by Dr. Bernard Weissman, University of North Carolina) were cultured in DMEM (Gibco) and supplemented with 10% TET free FBS (Corning) and 1% penicillin/streptomycin (Gibco). All cells were maintained at 37°C in a humidified incubator containing 5% CO_2_. All cell lines were routinely monitored for mycoplasma testing and STR profiled for cell line verification.

### The LSD1 screening biochemical assay

The LSD1 screening biochemical assay was performed as previously described (58). Briefly, the LSD1 biochemical kit was purchased from Cayman Chemical (Ann Arbor, MI). SP-2577 and SP-2513 were diluted to 20X the desired test concentration in 100% DMSO and 2.5 μL of the diluted drug sample was added to a black 384-well plate. The LSD1 enzyme stock was diluted 17-fold with assay buffer and 40 μL of the diluted LSD1 enzyme was added to the appropriate wells. Substrate, consisting of horseradish peroxidase, dimethyl K4 peptide corresponding to the first 21 amino acids of the N-terminal tail of histone H3, and 10-acetyl-3, 7-dihydroxyphenoxazine was then added to wells. Resorufin was analyzed on an Envision plate reader with an excitation wavelength of 530 nm and an emission wavelength of 595 nm.

### Cell Viability Assay

Cells were seeded in 96-well plates in triplicate at a density of 500 to 2,000 cells per well depending on the growth curve of each cell line. 24 h later, cells were treated with DMSO, LSD1 inhibitor SP-2577, or analog SP-2513 at increasing concentrations (0.001 to 10 μM). Cell viability was assessed with CellTiter-Glo (Promega) 72 h after treatment and IC_50_s were calculated using GraphPad Prism 8.0.

### Organoid Generation

SCCOHT cell lines (BIN67, SCCOHT-1, COV434) and the OCCC cell line TOV21G were seeded at 5000 cells per well in 96 well ultra-low attachment treated spheroid microplates (Corning) with the appropriate phenol red free medium containing 1.5% matrigel (Corning). Cells were maintained in culture for 72 h to generate spheroids for use in immune infiltration assays.

### Immune Infiltration Assay

#### SP-2577/SP-2513 Conditioned Media

The SCCOHT and OCCC cell lines were seeded in T-25 tissue culture treated flasks (Thermo Fisher) at 3 ×*10^5^* cells in 5mL of the appropriate phenol red free complete growth medium. After 48 h, at 70-80% confluency, the cells were treated with 3μM or 1μM of SP-2577 or SP-2513. After 72 h, the media was collected from the flasks and stored at −80°C until use in the infiltration assays.

#### Labeling PBMCs with RFP

Peripheral blood mononuclear cells (PBMC) (Lonza) were maintained in CTS OpTmizer T Cell expansion SFM (Thermo Fisher) in a T-75 suspension flask (Genesse Scientific) for 24 h at 37°C. After 24 h, cells were collected and washed with PBS (Gibco) and counted. Approximately 2.5 × *10*^*6*^ PBMCs were labeled with Molecular Probes Vybrant CM-Dil Cell Labeling Solution (RFP)(Invitrogen) by incubating the PBMCs with a 2 μM solution of CM-Dil for 45 min at 37°C in the dark and then for an additional 15 min at 4°C. After the incubation, cells were washed with PBS twice and resuspended in the appropriate complete growth medium.

#### Checkpoint blockade

The following monoclonal blocking antibodies (final concentration 10 μg/mL) were used for checkpoint blocking on tumors as well as T cells: functional grade PD-L1 (29E.2A3) and CTLA-4 (BN13) (Bioxcell, USA). In brief, 2 × 10^6^ PBMCs were treated with 10μg/mL of α-CTLA-4 antibody and incubated at 37°C incubator for 45 min on shaker. Then the cells were washed in serum free media and stained with Molecular Probes Vybrant CM-Dil Cell Labeling Solution as mentioned above. Alternatively, COV434 tumor organoids were incubated with 10 μg/mL of α-PD-L1 antibody and incubated at 37°C incubator for 1h. Then the media containing α-PD-L1 was removed carefully and fresh conditioned media was added on all organoids.

#### Immune infiltration and imaging

150μL of SP-2577 conditioned medium was added to each well containing a spheroid. A 5 μm HTS Transwell-96 Well permeable support receiver plate (Corning) was placed on each ultra-low attachment spheroid microplate (Corning) to allow for PBMC infiltration into the tumoroids. RFP-stained PBMCs were then seeded into inserts at 5 × 10^5^ cells/well to ensure tumoroid:PBMC cell ratio of 1:10. After 48 h, inserts were removed, and organoid microplates were analyzed by 3D Z-stack imaging and morphometric analysis with Cytation 5 software to quantify the lymphocyte infiltration.

### qPCR

COV434, BIN67, and SCOOHT-1 cells were seeded at a 1 × 10^6^ cells in 2 mL in 6-well tissue culture treated plates (Corning). After 24 hrs, cells were treated with 1 μM and 3 μM of SP-2577 as well as 3μM of SP-2513 for 72 h. DMSO was used as negative control. To quantify gene expression, total RNA was extracted (Qiagen RNeasy Mini Kit) and quantified by spectroscopy (Nanodrop ND-8000, Thermo Scientific). Samples were then reverse transcribed to cDNA using a high capacity cDNA reverse transcription kit (Applied Biosystems) and the MJ Research thermal cycler. cDNA was amplified, detected, and quantified using SYBR green reagents (Applied Biosystems) and the ViiA 7 Real-Time PCR System (Applied Biosystems). Data were normalized to GAPDH expression. List of primers used in this study are listed in Supplemental section.

### Fluorescent staining and imaging of Organoid

Organoids grown in Matrigel were initially fixed in 4% PFA for 1.5 h. After PBS washing, organoids were embedded in Histogel, processed with an automated tissue processor (Tissue-Tek VIP), and embedded into a paraffin block (Tissue-Tek TEC). Samples were sectioned at 4 μm onto poly-L-lysine coated slides and air-dried at room temperature over-night for any subsequent immunofluorescence staining. All slides for fluorescence were deparaffinized and antigen retrieved in pH 6 citrate buffer for a total of 40 min. After protein blocking, nuclei were stained with DAPI (Sigma). Infiltrating PBMCs were pre-stained with RFP as described previously. Imaging was performed on a Zeiss LSM880 fluorescent microscope with Zen Black software.

### Flow Cytometry and antibodies

A digestion step was performed for the harvest and characterization of lymphocytes infiltrated into the organoids. After immune infiltration experiments, organoids were removed from culture insert, washed with PBS twice to eliminate the PBMCs not inside the organoids and incubated with Gentle Cell Dissociation reagent (Stem Cell Technologies) for 2 min at RT and mechanically disrupted by pipetting. After complete disaggregation, single cell suspensions were dissolved in Cell Staining buffer (Biolegend, San Diego, California) and incubated at 4°C for 10 min prior antibody staining.

Cells were stained for CD45 (Miltenyi Biotech), CD3, CD8, CD4, CD56, and CD19 (Biolegend). Cell viability was assessed by negative live/dead antibody staining (Miltenyi Biotech). Analysis was performed on a BD FACS Canto II (BD Biosciences). BV421+ and single cells were gated out, followed by CD45+ cell selection. Analysis of CD3, CD8, CD4, CD56, and CD19 lymphocyte populations was performed with FlowJo software (Tree Star Inc.)

### Western Blot

COV434 pIND20 BRG1-2.7 cells were plated in 6 well tissue culture plates at a density of 5 × 10^5^ cells per well and left to adhere overnight. Once adherent, cells were treated with 1μM doxycycline (SIGMA) daily for 8 days. Cells were harvested daily and proteins were extracted from cell lysates and immunoblotted for SMARCA4 expression.

### TCGA analysis

TCGA PanCancer Atlas data was downloaded from cBioPortal. To identify SWI/SNF cancers cBioPortal was used to query samples with SMARCA4, ARID1A, SMARCB1, SMARCC1, and KDM1A mutations. Samples were removed that had only KDM1A mutations, KDM1A deletions, no profiling of SWI/SNF genes, or did not have expression data. Expression of KDM1A was plotted using RStudio.

### U-PLEX MSD analysis

MSD analysis was performed following manufacturer protocol. Briefly, conditioned media (CM) from SP-2577 and DMSO treated cells were collected and centrifuged at 1500 RPM for 5 min at 4°C to eliminate cell debris. CM was then concentrated using Centricon 10 KDa (Sigma) and 25 μL of the resulting CM was added in each well in the MSD plate (KIT #K15067L-2) and analyzed using Discovery Workbench 4.0 software.

### Statistical analysis

Student’s T-tests were performed using GraphPad Prism 8.0. Symbols for significance: NS, non-significant; *=p<0.05; **=p<0.01; ****=p<0.0001. All the *in vitro* experiments were performed in triplicate and repeated at least three times.

## Results

### LSD1 is highly expressed in SWI/SNF-mutant cancers and the LSD1 inhibitor SP-2577 inhibits SWI/SNF-mutation-dependent tumor cell proliferation

TGCA analysis using cBioPortal showed that LSD1 is highly expressed in the majority of human cancers, including in SWI/SNF-mutant tumors (59, 60) (Fig. 1 A and B). To determine the effects of SP-2577 on SWI/SNF-mutant cell viability, we performed drug-dose-response (DDR) studies with 72 h CellTiterGlo viability endpoints in SCCOHT (SMARCA4^™/−^), OCCC (ARID1A^™/−^), lung (SMARCA4^™/−^), kidney (SMARCB1^™/−^), and colorectal cancer cell lines (SMARCA4^™/−^). All cell lines were sensitive to the treatment and showed sub-micromolar IC_50_s (Table 1). SP-2513, an analog of SP-2577, which only poorly inhibits LSD1 enzymatic activity (Table 2), showed significantly higher 72 h IC_50_s that ranged from 3.5 to 10 μM (Table 1).

**Figure 1:**
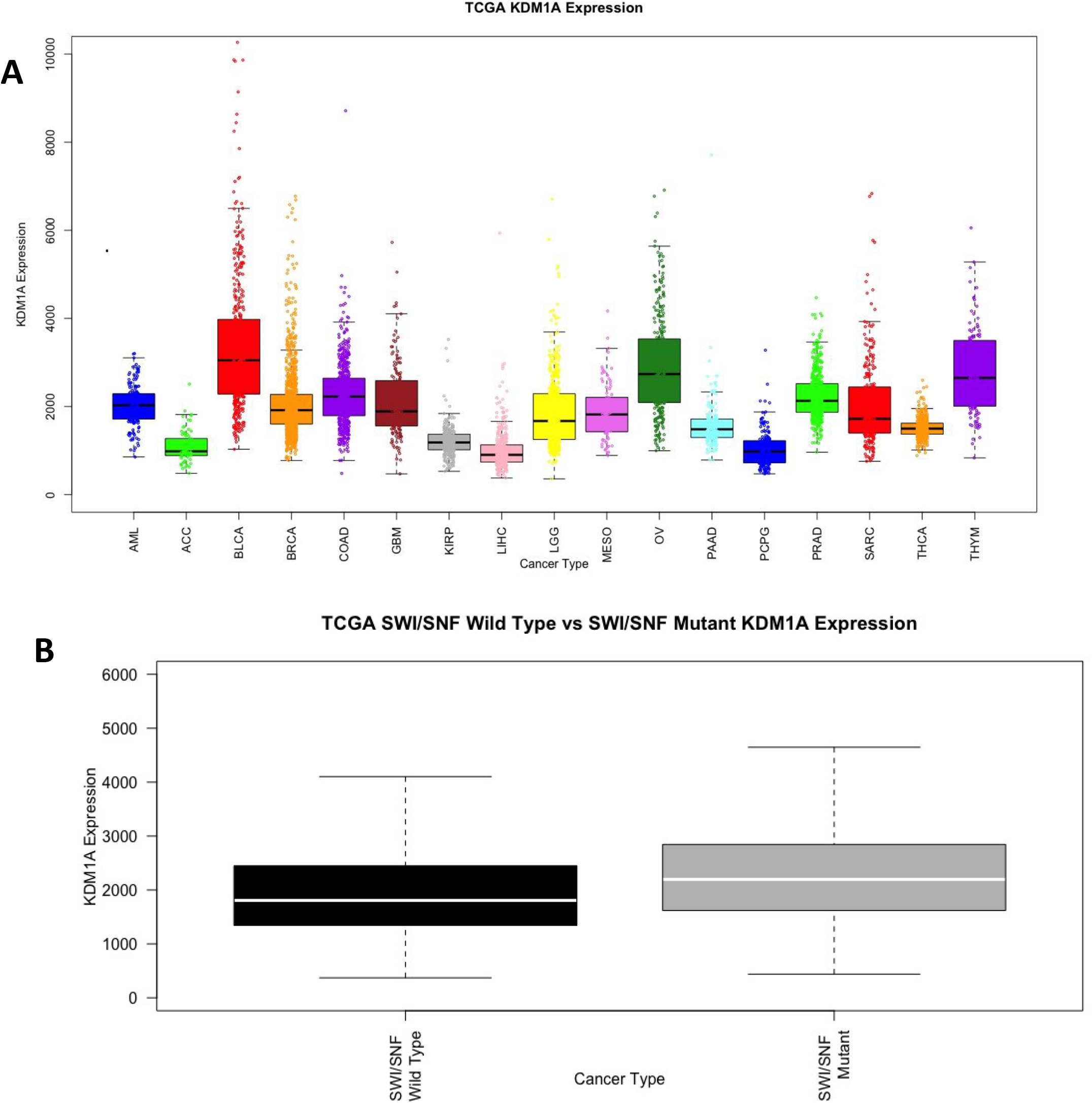
LSD1 expression in tumors and SP-2577 toxicity in SWI/SNF mutated tumors. (**A**) TCGA analysis for LSD1 expression in different human cancers. **(B)**TCGA analysis for LSD1 expression in SWI/SNF-mutated tumors.

**Table 1:** Cytotoxicity of SP-2577 and SP-2513 in SWI/SNF-mutated cancer cell lines.

| Table 1 |  |  |  |  |
| --- | --- | --- | --- | --- |
| Cell line | Cancer type | Drug IC<sub>50</sub> (μM) |  | SWI/SNF mutation |
|  |  | SP-2577 | SP-2513 |  |
| COV434 | SCCOHT | 0.529 | 9.985 | SMARCA4 |
| BIN67 | SCCOHT | 0.417 | 10 | SMARCA4 |
| SCCOHT-1 | SCCOHT | 1.098 | 3.65 | SMARCA4 |
| TOV21G | OCCC | 0.203 | 24.54 | ARID1A |
| SKOV3 | Ovary ADC | 0.013 | --- | SMARCC1 |
| A427 | Lung ADC | 0.1422 | --- | SMARCA4 |
| H522 | Lung ADC nscl | 2.819 | --- | SMARCA4 |
| A549 | Lung ADC | 0.248 | --- | SMARCA4 |
| H1299 | Lung ADC nscl | 0.212 | --- | SMARCA4 |
| G401 | Kidney Rhabdoid | 0.3874 | --- | SMARCB1 |
| G402 | Kidney renal leiomyoblast | 1.179 | --- | SMARCB1 |
| HCC15 | CRC | 0.117 | --- | SMARCA4 |

**Table 2:** Inhibition of LSD1 enzymatic activity of SP-2577 and SP-2513.

| Table 2 |  |
| --- | --- |
| Drug | IC<sub>50</sub> (μM) |
| SP-2577 | 0.013 |
| SP-2513 | >1 |

LSD1 screening biochemical assay

### Inhibition of LSD1 by SP-2577 promotes ERV expression and induces expression and release of effector T cell attracting chemokines in conditioned medium

Inhibition of LSD1 activity has been suggested to enhance anti-tumor immunity through activation of ERVs and production of dsRNA, followed by activation of an IFN-β-dependent immune response (51). To investigate whether SP-2577 treatment could promote a similar response in SCCOHT tumors, we analyzed the expression of ERVL, HERVK, and IFN-β in SCCOHT cell lines (COV434, BIN 67 and SCCOHT-1) treated with SP-2577. After 72 h treatment with 3μM SP-2577, quantitative PCR (qPCR) analysis showed that SP-2577 significantly upregulated the expression of ERVs, IFN-β, and interferon-stimulated genes ISG15 and CXCL10, suggesting activation of an IFN-dependent immune response (Fig. 2A). Recently it has been shown that LSD1 inhibition in triple negative breast cancer cell lines induces expression of CD8^+^ T cell-attracting chemokines including CCL5, CXCL9, and CXCL10 (61). The expression of these genes along with PD-L1 was shown to have increased H3K4me2 levels at proximal promoter regions (61) in response to LSD1 inhibition. To determine whether SCCOHT cell lines release CD8^+^ T cell-attracting chemokines after LSD1 inhibition, we analyzed the conditioned media from COV434 cells treated with SP-2577 for 72 h. Our U-PLEX MSD data showed that treatment with SP-2577 stimulated chemokine secretion in COV434 culture medium (Fig. 2B). Further, secretion of cytokines including IL-1**β**, IL-2, and IL-8 were also observed in the treated conditioned medium. Together, these data suggest that SP-2577 may play a role in the promotion of anti-tumor immunity.

**Figure 2:**
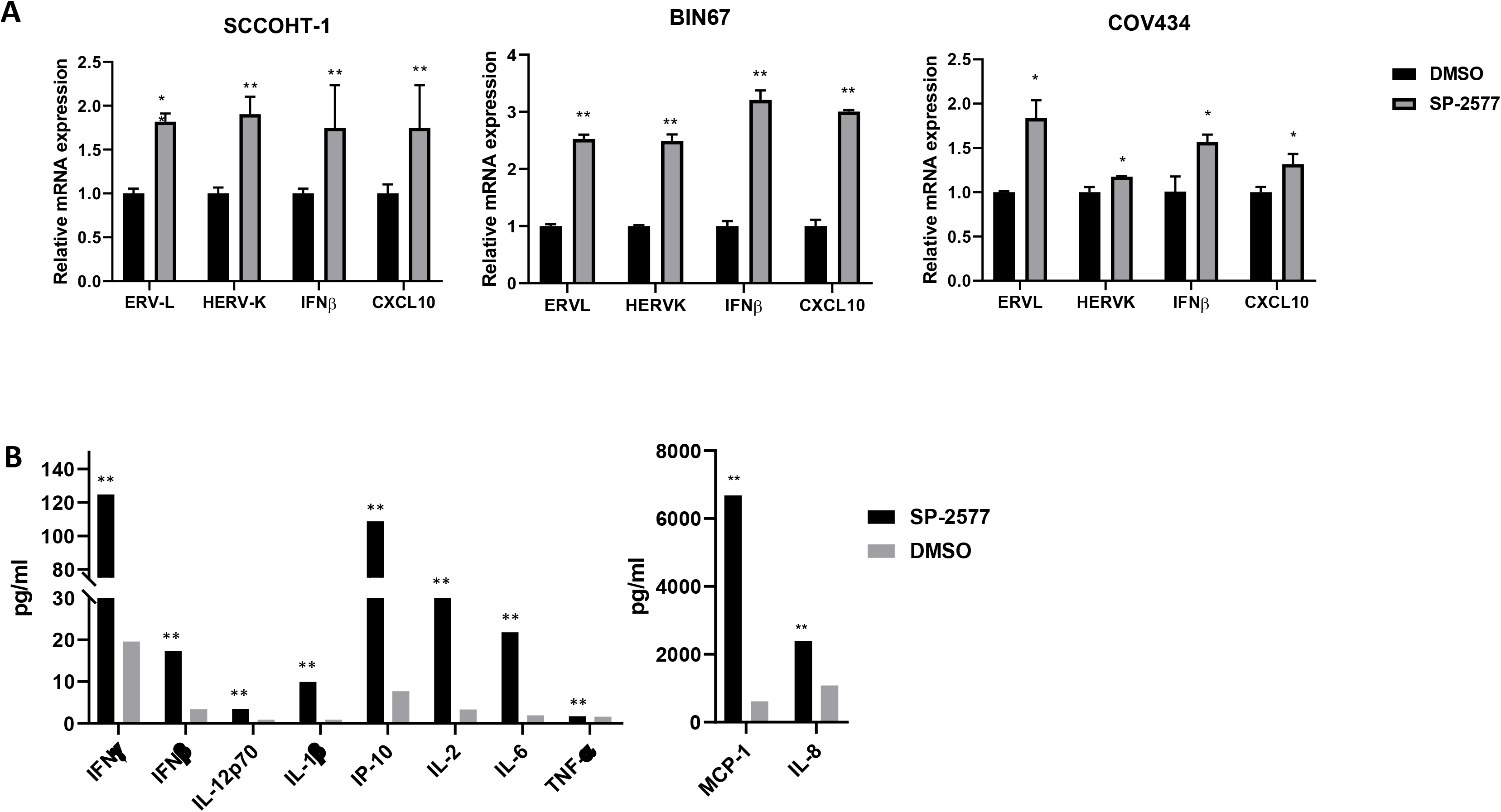
SP-2577 promotes ERVs expression and activation of IFNβ pathway in SCCOHT cell lines. (**A**) qPCR analysis of SCCOHT cell lines BIN67, COV434 and SCCOHT-1 after 72h of SP-2577 treatment showing increased expression of ERVs and IFN pathway cytokines. (**B**) MSD panel of chemokines and cytokines from SCCOHT COV434 cell conditioned media in the presence or absence of SP-2577 for 72h. *=p<0.05, **=p<0.01.

### SP-2577 promotes lymphocyte infiltration in 3D culture of SCCOHT cell lines

To examine whether SP-2577-dependent cytokine and chemokine secretion could enhance lymphocyte trafficking and tumor infiltration, we carried out an *ex vivo* migration/infiltration assay. SCCOHT cell lines COV434, BIN67, and SCCOHT-1 were grown in low concentration matrigel to form 3D organoids. To prevent SP-2577 cytotoxic damage from directly impacting cell viability, the organoids were cultured in conditioned media from SCCOHT cell lines collected 72 h post-treatment with SP-2577. At this time point there was no drug present in the conditioned medium. Migration and infiltration of lymphocytes was determined utilizing RFP-stained human allogeneic PBMCs. PBMCs were added to organoid cultures and migration toward the organoids was assessed by three dimensional Z-stack imaging and morphometric analysis (Cytation 5, BIOTECK). SP-2577 promoted PBMC infiltration in organoids more efficiently than the less active analog SP-2513 (Fig. 3A). Immunofluorescence analysis further demonstrated the presence of stained PBMCs in the sectioned organoids that were treated with SP-2577 conditioned medium while they were absent in organoids cultured in SP-2513 or DMSO-conditioned medium (Fig. 3B). Finally, to determine the dependency of PBMC infiltration on SP-2577, we performed *ex vivo* migration/infiltration studies with higher concentrations of the LSD1 inhibitor (0.03 −3μM). SP-2577 induced PBMC infiltration into the SCCOHT organoids in a dose-dependent manner (Fig. 3C). Together, these observations suggest that the chemokines and cytokines secreted by SCCOHT cells in response to SP-2577 treatment promote the migration and infiltration of PBMCs in tumor organoids.

**Figure 3:**
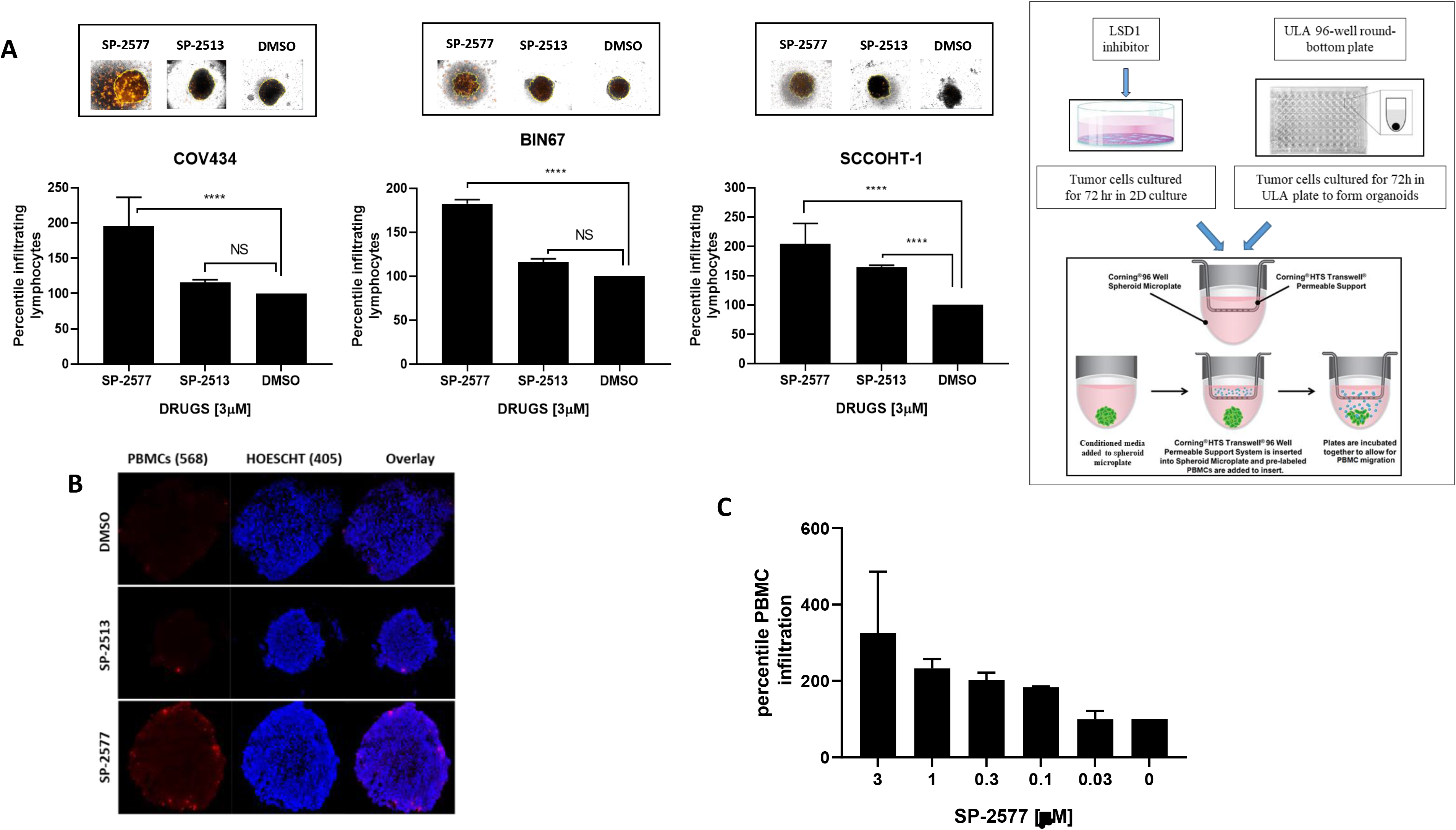
SP-2577 promotes lymphocytes infiltration in SCCOHT tumor organoids. (**A**) Immune infiltration assay in SCCOHT organoids imaging analysis. COV434, BIN67 and SCCOHT-1 derived-organoids were incubated with conditioned medium pretreated with 3 μM SP-2577, SP-2513 or DMSO in the presence of RFP-tagged PBMCs. After 48h the levels of lymphocyte infiltration were measured by z-stack analysis by Cytation 5 imaging. P values for COV434=<0.0001, BIN67 = 0.0016, and SCCOHT-1 = 0.0196. Right panel: Experimental design (**B**) IF analysis: RFP-tagged lymphocytes infiltration in SCCOHT organoids micro sections was assessed by analysis of RFP levels on confocal scope after 48h co-culture in presence of SP-2577 or SP-2513 conditioned medium. (**C**) Immune infiltration assay of COV434 cells treated with increased concentration of SP-2577 for 48h. The levels of lymphocytes infiltration correlate with SP-2577 dose.

### Treatment with SP-2577 promotes infiltration of T cells CD8+ into SCCOHT organoids

Next we investigated which lymphocyte populations infiltrated the in SCCOHT organoids after SP-2577 treatment *in vitro*. Flow cytometry analysis of the dissociated COV434 organoids shown the predominance of CD3^+^ T lineage cells (Fig. 4 Ai, Bi). CD8^+^ T cells percentile in SP-2577 treated organoids is highly significant compared to the untreated (Fig. 4 Aii, Bii). Similarly, CD4^+^ and CD4^+^CD8^+^ Double Positive T cells levels increased significantly after treatment with SP-2577 (Fig. 4 Aii, Bii). CD56^+^ NKT cells and CD19^+^ B cells were present in small percentile in the total CD45^+^ cell population. CD56^+^ NKT cells level increased approximately 15 folds after treatment with SP-2577 (Fig. 4 Aiii, Biii), while CD19^+^ B cells decreased by approximately 3 folds (Fig. 4 Aiv, Biv). All together these data suggest that treatment with SP-2577 promotes T cells and NKT cells infiltration into the tumor which leads to tumor cytotoxicity.

**Figure 4:**
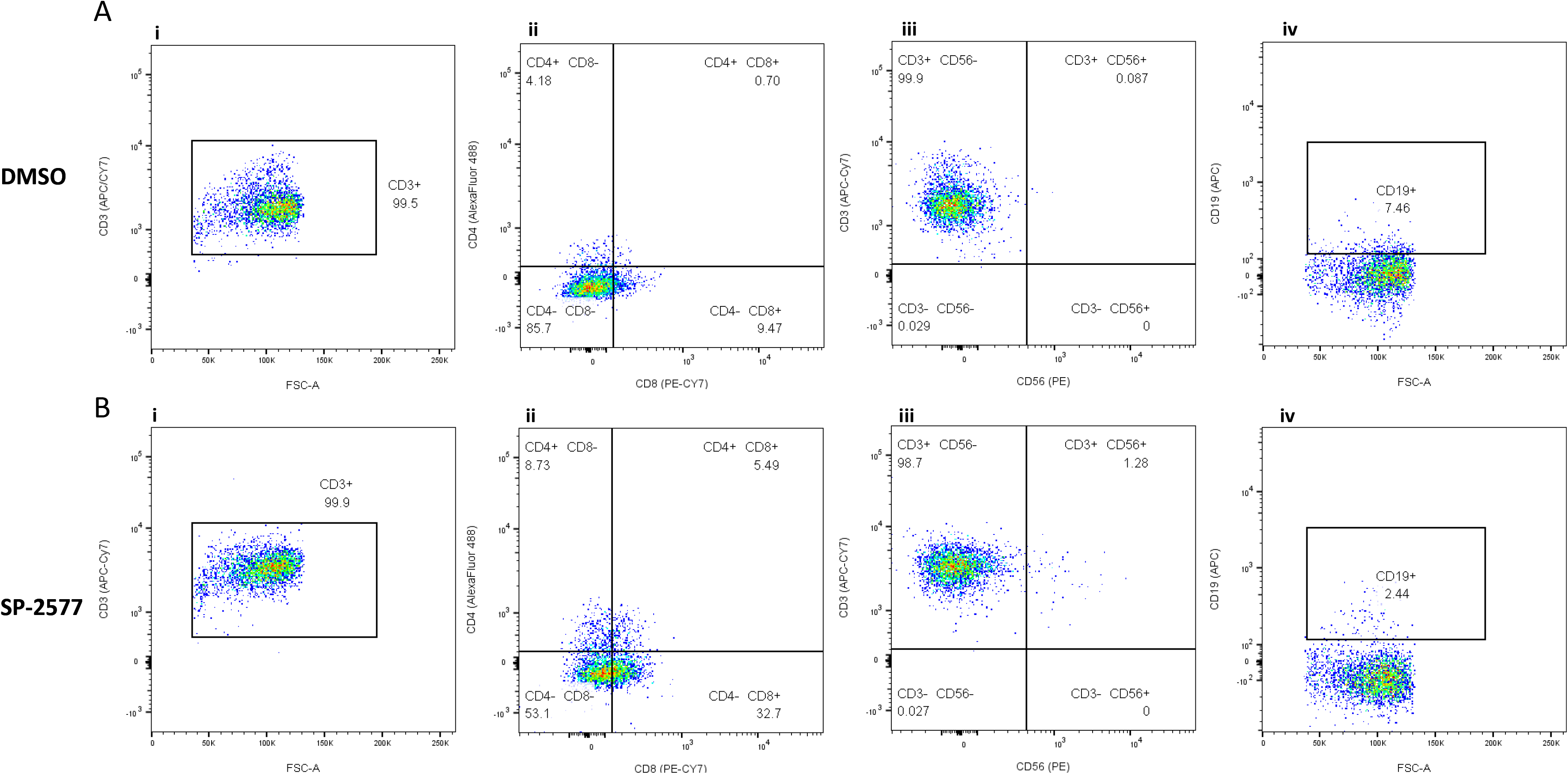
Flow cytometry analysis of the COV434 organoids after lymphocyte infiltration assay. COV434 organoids were dissociated and the infiltrated lymphocytes were stained with different markers and analyzed in Flow Cytometry as described in materials and methods. Panel A and B are showing the status of lymphocytes infiltrated in DMSO control organoids and SP-2577 treated organoids respectively in the representative dot plots. **(Ai and Bi)**The dot plots showed the CD3+ population in CD45+ gated lymphocytes. **(Aii and Bii)**The status of CD4+ and CD8+ T cells gated on CD3+ cells are presented in different quadrants. **(Aiii and Biii)**Percentage of CD56+ NK in CD3+ cells are presented with quadrant statistics. **(Aiv and Biv)**The status of CD19+ B cells are shown in CD3+ cells.

### Inhibition of LSD1 by SP-2577 induces expression of PD-L1

LSD1 expression has been shown to negatively correlate with expression of immune-related genes including PD-L1 (61). Further, LSD1 has been shown to play a critical role in epigenetic silencing of the expression of PD-1 (62) and LSD1 inhibition results in increased expression of PD-L1 in tumor cells (51, 61). To investigate the effect of SP-2577-dependent LSD1 inhibition on the expression of PD-L1 in SCCOHT, we treated COV434 organoids with 3 μM SP-2577 and performed qPCR analysis for PD-L1 expression. We observed a significant increase in PD-L1 expression (Fig. 5A). Next, we tested if checkpoint blockade could amplify the immune cell infiltration effect in the presence of SP-2577 in immune-organoids. As shown in Fig. 5B, co-treatment of low doses (100 nM) of SP-2577 with α-PD-L1 significantly increased lymphocyte infiltration. Moreover, the combination of α-PD-L1 and α-CTLA-4 was able to significantly enhance the infiltration of PBMCs in the SP-2577-treated COV434 organoids (Fig. 5B).

**Figure 5:**
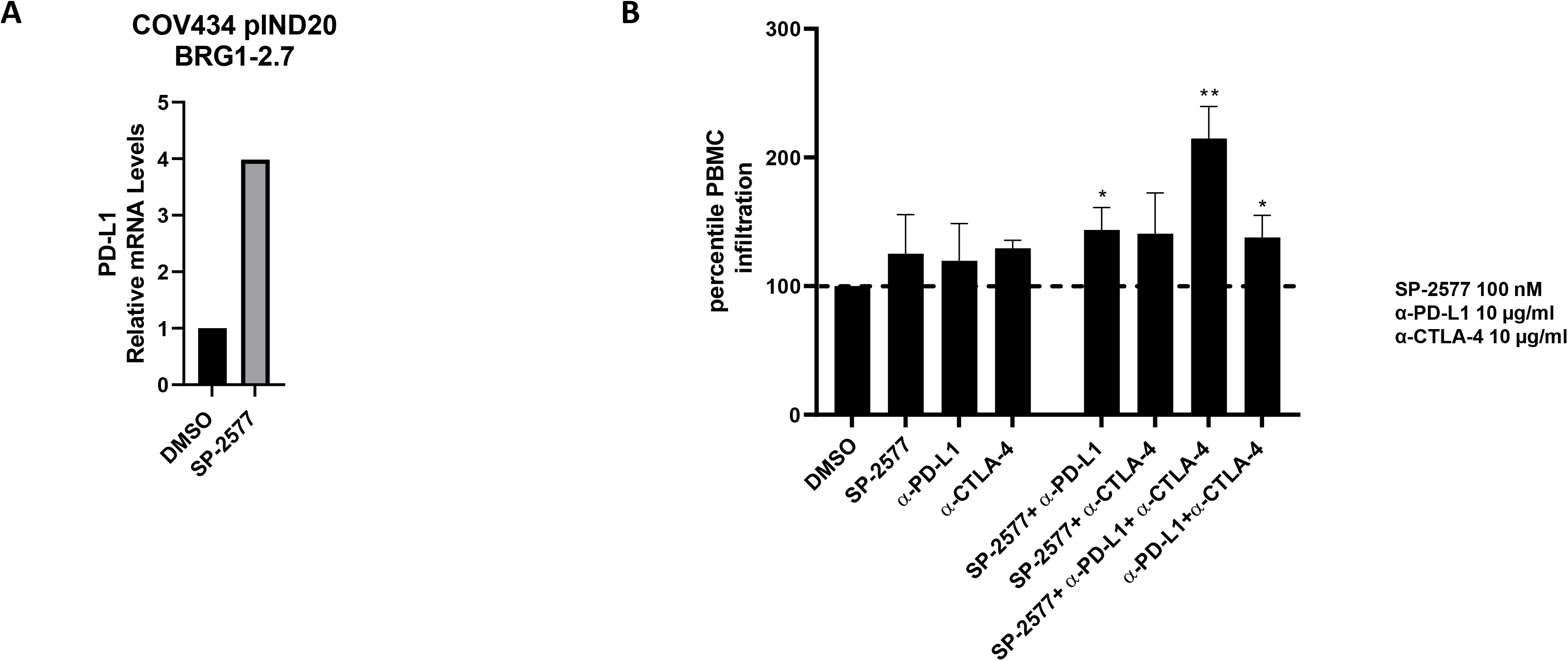
SP-2577 promotes PD-L1 expression and rescues checkpoint inhibition sensitivity in SCCOHT COV 434 cell line. (**A**) RT-PCR analysis of COV434 cells after SP-2577 treatment shows increase of PD-L1 expression levels, which are lost after Doxycycline treatment. (**B**) Co-treatment of SCCOHT organoids with SP-2577 and anti PD-L1 antibodies and lymphocytes treated with anti CTLA-4 antibodies showed an increase in lymphocyte infiltration as checked after 48h.

### SMARCA4 re-expression in SCCOHT cell lines blocks lymphocyte infiltration

Although SCCOHT is characterized by low tumor mutation burden, recent studies have indicated that the tumor microenvironment of SCCOHT is similar to other immunogenic tumors that respond to checkpoint blockade (37, 40, 63), suggesting that mutations in the subunits of the SWI/SNF complex may contribute to immunotherapy sensitivity. We questioned whether the restoration of SWI/SNF functionality in SCCOHT would affect the lymphocyte trafficking and tumor infiltration observed with SP-2577 treatment. To investigate this hypothesis, we used an isogenic COV434 cell line (COV434 pIND20 BRG1 2.7) where SMARCA4 (*BRG1*) was re-expressed under a doxycycline inducible system (29). qPCR and Western blotting analysis confirmed the expression of SMARCA4 in doxycycline-treated cells (Fig. 6A, B). SMARCA2 (*BRM*), which is normally silent in SCCOHT tumors, was also overexpressed by 14-fold after BRG1 induction (Fig. 6B). We observed a significant reduction in the level of infiltrated lymphocytes in SMARCA4 re-expressed organoids after SP-2577 treatment (Fig. 6C). Remarkably, expression of ERVL, HERVK, IFNβ, as well as interferon-stimulated genes ISG15 and CXCL10 was significantly downregulated in SMARCA4-induced cells after treatment with SP-2577 (Fig. 6D). In addition, re-expression of SMARCA4 resulted in downregulation of PD-L1 expression in SCCOHT cell lines (Fig. 6E). Lastly, the production and secretion of cytokines and chemokines that were induced by SP-2577 treatment were negatively affected by the re-expression of SMARCA4 in SCCOHT cells (Fig. 6F).

**Figure 6:**
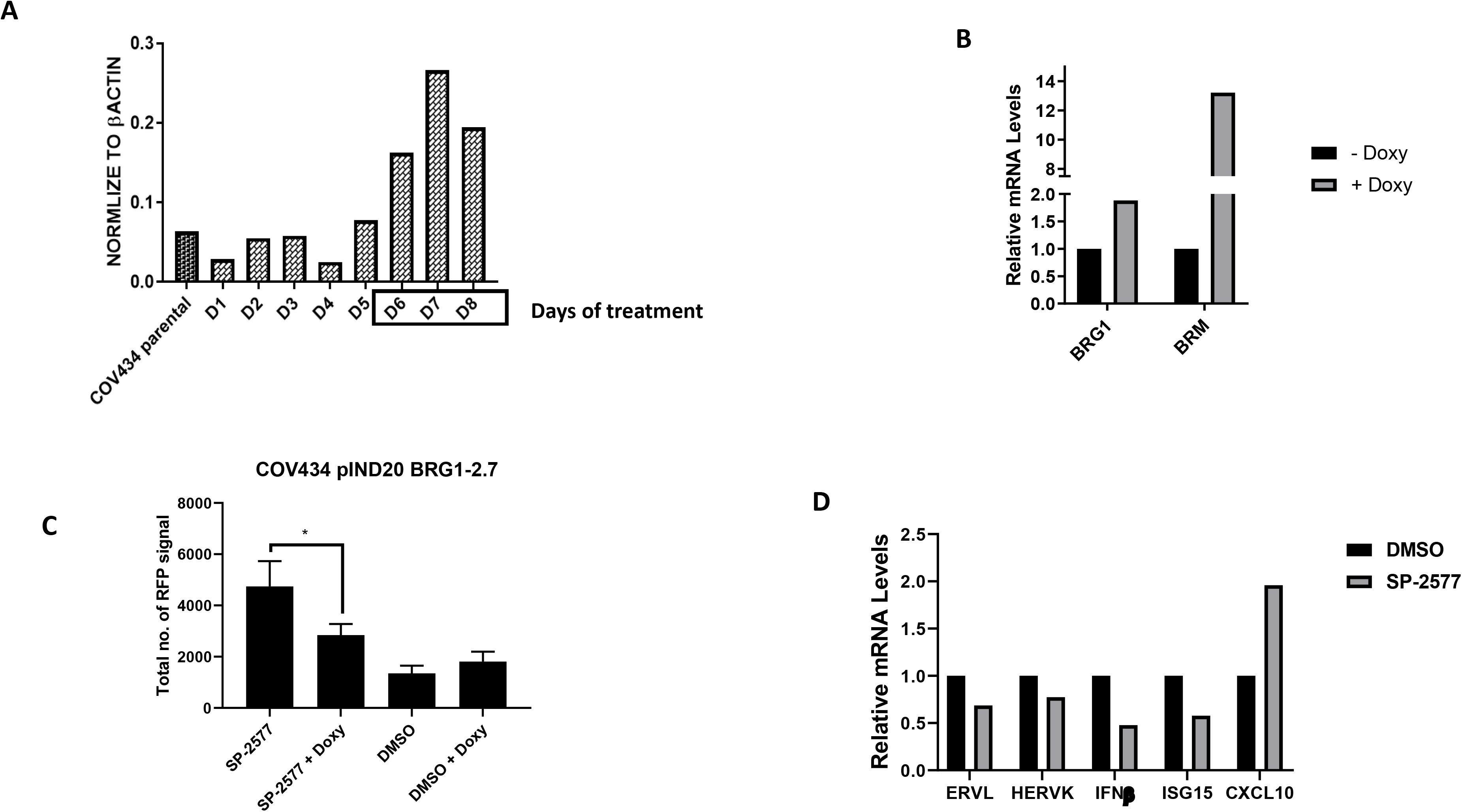
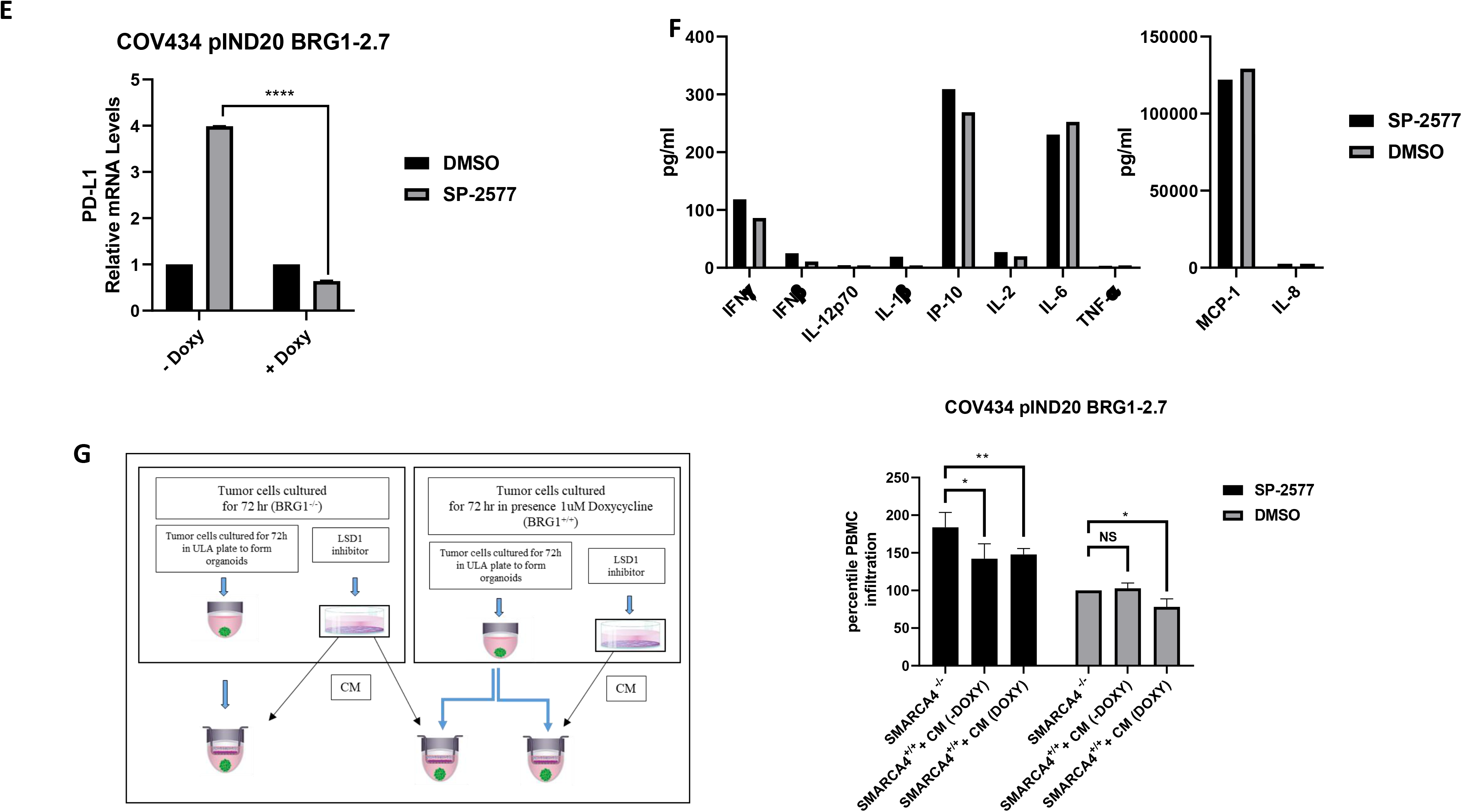
SMARCA4 re-expression in SCCOHT cell lines blocks lymphocyte infiltration. Densitometry analysis of western blot for SMARCA4 expression level in COV434 pIND20 BRG1-2.7 after 1μM Doxycycline daily treatment (D=days) (**B**) RT-PCR for BRG1 and BRM expression levels in COV434 pIND20 BRG1-2.7. Both gene expression increased after SP-2577 treatment. (**C**) The levels of lymphocyte infiltration are significantly reduced after SMARCA4 re-expression in COV434 pIND20 BRG1-2.7 (P value = 0.038). (**D**) RT-PCR analysis of SMARCA4-induced COV434 pIND20 BRG1-2.7 after 72h SP2577 treatment shows decrease of ERVs activity and INF expression. (**E**) RT-PCR analysis of COV434 cells after SP-2577 treatment shows increase of PD-L1 expression levels, which are lost after Doxycycline treatment **(F)**MSD analysis of SMARCA4-induced COV434 pIND20 BRG1-2.7 after 72h SP-2577 treatment shows significant decrease of cyto/chemokines released in medium after treatment. (**G**) Treatment of SMARCA4-induced COV434 pIND20 BRG1-2.7 organoids with conditioned medium (CM) from SMARCA4-deficient COV434 pIND20 BRG1-2.7 does not affect lymphocyte infiltration suggesting SMARCA4 plays role in immune response. Left panel: Experimental design.

To determine if the reduction of lymphocyte infiltration was solely dependent on the re-expression of SMARCA4, we cultured doxycycline treated COV434 pIND20 BRG1 2.7 organoids with conditioned medium generated from SP-2577-treated parental SCCOHT cells lacking SMARCA4 expression. As shown in Fig. 6G, the presence of cytokines and chemokines in SP-2577-treated conditioned medium was not sufficient to overcome the impaired infiltration of lymphocytes in SMARCA4-expressing organoids, suggesting that SMARCA4-dependent epigenetic changes may have altered the immunogenicity of the organoids.

### SP-2577 promotes lymphocyte infiltration in ARID1A deficient cells

To investigate if SP-2577 promotes tumor immune response in other SWI/SNF-mutant tumor types, we performed *ex vivo* migration and infiltration of lymphocytes in ARID1A-deficient OCCC cells. These studies were conducted in isogenic TOV21G cells, which re-express ARID1A under the control of doxycycline treatment. As previously observed in SCCOHT cell lines, SP-2577 promotes lymphocyte infiltration in a dose-dependent manner in parental TOV21G organoids (Fig. 7A). Re-expression of ARID1A resulted in a significant reduction in infiltration of lymphocytes (Fig. 7B). In addition, the combination of α-PD-L1 and α-CTLA-4 was able to significantly enhance the infiltration of PBMCs in the low dose (300 nM) SP-2577-treated parental TOV21G organoids (Fig. 7C).

**Figure 7:**
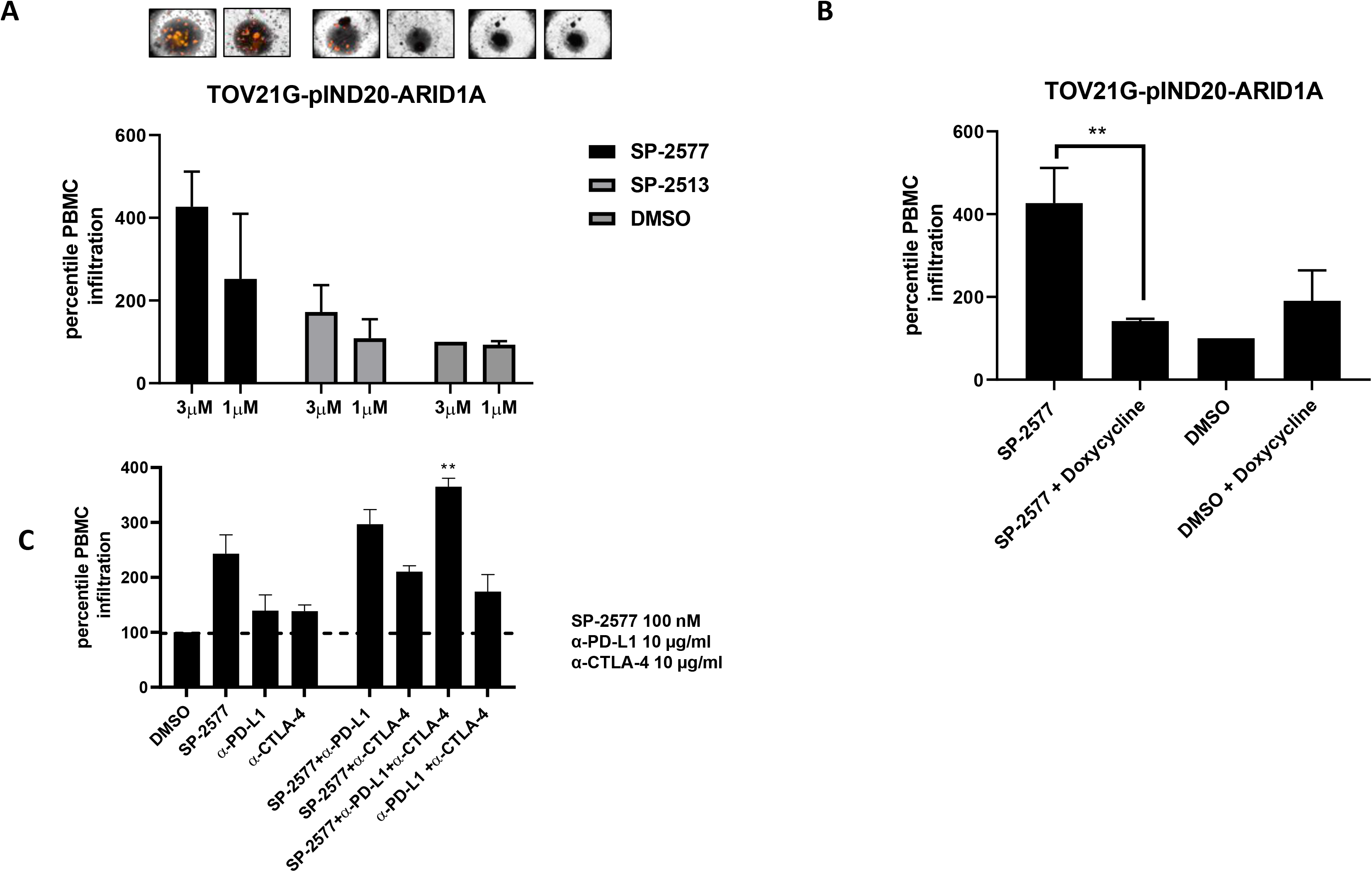
SP-2577 promotes lymphocyte infiltration in ARID1A deficient cells. (**A**) level of lymphocytes infiltration is dependent of SP-2577 treatment in TOV21G pIND20-ARID1A cells line. (**B**) lymphocytes infiltration is significantly reduced in TOV21G pIND20-ARID1A cells after ARID1A re-expression upon Doxycycline treatment (P value = 0.004). (**C**) co-treatment of SP-2577 and CTLA-4 antibodies in the immune infiltration assay showed an increase in lymphocyte infiltration in TOV21G-pIND20-ARID1A after 48h.

## Discussion

SWI/SNF complexes have previously been implicated in regulation of the immune system, particularly in enhancing interferon-stimulated gene (ISG) expression (64) through a STAT-dependent mechanism. However, it has been reported that inactivating mutations in SWI/SNF subunits (ARID1A, PBRM1, SMARCB1, and SMARCA4) also sensitize cancer cells to T cell-mediated destruction (37). SWI/SNF loss-of-function enhances the expression of immune checkpoint regulators and neoantigen presentation (37, 39, 65). Even monogenic tumors like SCCOHTs are immunogenic and exhibit biologically significant levels of T cell infiltration and PD-L1 expression (40). It has been reported that several patients with mutations in the SWI/SNF complex have benefited from checkpoint blockade immunotherapy (37, 39, 40). Although immune checkpoint inhibitor treatment has shown promising results, only a minority of treated patients exhibit durable responses (40, 66). Additionally, many of those who initially respond to treatment eventually experience relapse due to acquired resistance. As with conventional cancer therapies, one way to improve clinical responses with immune checkpoint blockade is through combination therapy strategies.

Numerous studies have revealed that epigenetic modulation plays a key role in tumor immune escape. Cancer cells show frequent loss or epigenetic silencing of the cytosolic DNA sensor cGAS and/or STING to promote immune evasion (67). Conversely, aberrant LSD1 activity suppresses the expression of immune protective factors (61) such as PD-L1 and CTLA-4. Recently, it has been shown that inhibition of LSD1 significantly increases tumor immunogenicity (51) primarily through changes in methylation levels. An increased methylation level promotes endogenous retroviral (ERV) expression, leading to dsRNA stress. Methylation of the RISC complex member AGO2 results in destabilization of the integrity of complex and inhibition of dsRNA degradation. dsRNA stress activates IFN-dependent immune response that consequently sensitizes tumors to T cell immunity and T cell infiltration. As LSD1 is highly expressed in SCCOHT cell lines and interacts with members of the SWI/SNF complex to regulate gene expression (55, 68, 69), it is an ideal therapeutic target for the treatment of SCCOHT and potentially other SWI/SNF loss of function-dependent cancers.

In this study, we demonstrate that inhibition of LSD1 activity by the reversible inhibitor SP-2577 (Seclidemstat) induces the expression of ERVs and IFN**β** in SCCOHT. In agreement with the previous observation that LSD1 negatively regulates expression of chemokines and immune protective factors such as PD-L1 (61), inhibition of LSD1 with SP-2577 promoted expression of chemokines and PD-L1 in SCCOHT cell lines, which results in activation of T cell infiltration.

The contribution of ERVs to the immune response is not limited to the dsRNA/MDA5 interaction followed by activation of the IFN pathway. Tumor neoantigen analysis has shown that ERVs can encode for strictly tumor-specific antigens, otherwise silent in normal tissue, capable of eliciting T cell specific antitumor immunity. Schiavetti et al have shown that ERV-K antigens are highly expressed in a variety of malignancies such as breast, melanoma, sarcoma, lymphoma, and bladder cancer (70), and the ERV-E encoded antigen is selectively expressed in RCC kidney tumors (71). Moreover, it has been shown that in epithelial ovarian cancer and in colon cancer high levels of ERVs expression correlates robustly to immune checkpoint therapy response (72, 73). Expression of ERV elements is subjected to genome-wide regulation by epigenetic silencing (74, 75). However, many ERVs are still transcribed in adult cells and contribute to autoimmune pathologies such as systemic lupus erythematous and Aicardi–Goutières syndrome (76). Similarly, dysregulation of epigenetic pathways also contributes to reactivation of ERV elements in tumor cells (77).

SWI/SNF may play a role in the establishment of ERV silencing. In embryonic stem cells SMARCAD1, a SWI/SNF-like chromatin remodeler, negatively regulates the retrotransposon activity through recruitment of KRAB associated protein 1 (KAP1) (78). Docking KAP1 at the ERV elements in the genome triggers the formation of a complex with the histone methyltransferase SETDB1, resulting in formation of H3K9me3 and silencing of ERV class I and II. Lack of SMARCAD1 compromises the stability of KAP1-SETDB1 association at ERVs, which lead to reduction of H3K9me3 and activation of ERV transcription. Restoration of SMARCAD1 activity reverses the ERV upregulation (78). Similarly, SWI/SNF complex members, including SMARCA4 and LSH1, are known to ensure silencing of retrotransposons in the stem cells through interaction with DNMT3a and HDACs (75, 78). Transcriptome analysis of rhabdoid tumors cell lines with inducible SMARCB1 re-expression identified significant ERVs overexpression in SMARCB1-deficient conditions (79). In support of this, our studies show that re-expression of SMARCA4 in SCCOHT or ARID1A in OCCC results in the loss of ERV expression and reduction of T cell infiltration, even in the presence of conditioned medium enriched in cytokines and chemokines, implying that T cell infiltration may be related to the loss of SWI/SNF complex function. Together these results suggest that SWI/SNF deficiency plays a crucial role on ERVs epigenetic silencing, resulting in enhancement of tumor immunogenicity. Moreover, LSD1-dependent histone modifications result in significant downregulation of ERVs (74, 80). By inhibiting LSD1, SP-2577 ensures increased ERV activation and cytokine production, resulting in enhanced T cell immune response. In addition, SP-2577 treatment promotes PD-L1 expression, and co-treatment with α-PD-L1 or α-CTLA-4 antibodies significantly amplifies the CD8^+^ T cell infiltration in SCCOHT and OCCC immune organoids.

Collectively, these findings demonstrate the important role of SP-2577 in the regulation of T cell recruitment to the tumor microenvironment and highlight the potential of combining SP-2577 with checkpoint immunotherapy in SCCOHT, OCCC, and other SWI/SNF mutated malignancies.

## Author contributions

RS, TGH, AW and RRdV conducted the experiments, analyzed the results, and wrote most of the paper. KD performed the TGCA analysis. HV, MRK and WPDH provided advice on experimental design, interpretation of results and preparation of the manuscript. TT, RL, SHS, SDA, JL, BW and JMT provided advice on the preparation of the manuscript. SS conceived the idea for the project, provided advice on experimental design, interpretation of results and preparation of the manuscript.

